# Clinical *Campylobacter jejuni* isolates: genomes and genetic tools

**DOI:** 10.64898/2026.05.21.726778

**Authors:** Vida Nasrollahi, Gary W. Foo, Tahani Jaafar, Abdelbaset A. Elzagallaai, Michael J. Rieder, Bogumil J. Karas

## Abstract

*Campylobacter jejuni* is a major cause of food-borne gastroenteritis and is responsible for substantial mortality and economic losses in meat and dairy production. Detecting *C. jejuni* in contaminated food samples remains difficult because current assays are culture-based, slow, and can yield false positives. As a result, contamination may not be identified for several days, limiting detection at the point of production. Developing improved assays has also been challenging because *Campylobacter* genetics and the biology of clinical isolates remain poorly understood. Here, we expand the *C. jejuni* genetic toolbox by sequencing two strains, HC1 and RM1164, derived from patient and food samples. We identified two cryptic plasmids in HC1, one potentially capable of conjugation and another conferring tetracycline resistance. We also engineered a mobilizable plasmid carrying an *OriT* sequence that can be transferred from *Escherichia coli* donor strains to *C. jejuni* RM1164 by conjugation. Together, these clinical isolates and the plasmid system expand the genetic tools available for *C. jejuni*.

## Introduction

Food-borne illnesses are a growing public health concern, with cases expected to rise across North America [1,2]. Millions of infections are reported each year, leading to substantial mortality and billions of dollars in food recall costs [1,3]. Pathogenic bacteria are the main causes of food-borne disease, and *Campylobacter jejuni* is consistently ranked as one of the most common bacterial causes of gastroenteritis [4]. Infections are often linked to raw or undercooked poultry, meat, or other contaminated dairy produce [5]. In addition to acute gastroenteritis, campylobacteriosis can trigger severe autoimmune sequelae, including Guillain–Barré and Miller–Fisher syndromes [6,7]. Cases of campylobacteriosis are expected to rise in coming years, as climate change may promote conditions favourable for Campylobacter growth and extend its seasonal prevalence in avian hosts [1,8]. Together, these factors highlight the importance of improving *Campylobacter* detection in contaminated food products and reducing its public health burden.

The average *C. jejuni* genome is between 1.61 and 1.87 Mb, with a GC content of ∼30.9% and 1600-1800 protein coding genes [9,10]. Comparative genomics indicates a conserved core genome comprised of approximately 1400-1500 core genes [11]. Some *C. jejuni* strains also harbour cryptic plasmids that carry antimicrobial resistance genes and can be self-transmissible, such as pTet and pVir, which span 1.3–208 kb and encode distinct replicon types and relaxases that may contribute to strain-specific pathogenicity in *C. jejuni* [12]. Type III (T3SS) and type VI (T6SS) secretion systems have been described in *C. jejuni*, with about 16–20% of sequenced strains carrying T6SS loci associated with adhesion and invasion. Strains that encode T3SS express *Campylobacter* invasion antigens that promote uptake of *C. jejuni* into host cells [13].

A genetic toolbox for *C. jejuni* is currently under development, containing several methods for DNA deliver, including natural transformation, electroporation, and conjugation [14]. Natural transformation into *C. jejuni* can be achieved by spreading a layer of culture across an agar plate and adding an additional layer of transformant DNA carrying a selection marker [15]. Electroporation into *C. jejuni* has also been previously documented, including protocols and settings for DNA transfer and producing electrocompetent *C. jejuni* cells [16]. For conjugation, the mobilizable plasmid pRK212.1 has been used to deliver replicating vectors into *C. jejuni* [14] .Other shuttle vectors have also been developed that carry *Campylobacter* and *E. coli* replicons, multiple cloning sites, and selectable markers such as kanamycin and ampicillin [17]. There are also methods for genome mutagenesis that use Tn5, EZ:Tn5, and mariner transposons [18–20]. Together, these tools enable gene knockouts, chromosomal insertions, random mutagenesis, and recombinant gene expression in *C. jejuni*. However, we and others have found that many of these approaches perform inconsistently, with transformation efficiencies varying widely between *C. jejuni* strains [21,22]. Most of these tools were developed for commonly used reference strains, including NCTC 11168, and have not been well documented in more virulent isolates such as C. jejuni 81-17623 [23]. This highlights the need for *C. jejuni* genetic tools that work more broadly, especially in clinically relevant isolates.

Here, we build on the existing *Campylobacter* genetic toolbox by introducing two new *C. jejuni* strains, RM1164 and HC1, isolated from contaminated food and clinical samples. We sequenced both strains to identify genomic features of interest and to assess the presence of cryptic plasmids. Two cryptic plasmids were identified in *C. jejuni* strain HC1 that appear to carry conjugation machinery, virulence factors, and genes conferring resistance to tetracycline. To host recombinant genes in *C. jejuni*, we developed a shuttle plasmid (pCJ01) capable of replication in *E. coli* and *C. jejuni*. To facilitate genetic manipulation of *C. jejuni*, we developed a shuttle plasmid (pCJ01) capable of replication in both *E. coli* and *C. jejuni*. pCJ01 also carries an origin of transfer, allowing its mobilization from a non-pathogenic *E. coli* donor strain into *C. jejuni*. Together, this work expands the *Campylobacter* genetic toolbox by introducing two diverse *C. jejuni* isolates and demonstrating plasmid transfer by conjugation.

## Materials and methods

### Microbial strains and growth conditions

*Campylobacter jejuni* strain RM1164 was provided by Dr. Michael Rieder. *C. jejuni* strain Human Clinical Isolate 1 (HC1) was provided by London Health Sciences Centre (LHSC). *C. jejuni* strains were grown in a microaerobic atmosphere (85% N_2_, 10% CO_2_, and 5% O_2_) at 37°C on Columbia Blood Agar Base (CBA) (Difco^™^279240) supplemented with 5% sheep blood (Cedarlane Laboratories Ltd., Hornby, ON, Canada), 10 *µ*g/mL vancomycin, and 5 *µ*g/mL trimethoprim as background antibiotics. Growth media was supplemented with 30 *µ*g/mL kanamycin as required. *Saccharomyces cerevisiae* strain VL6-48 (American Type Culture Collection [ATCC], catalog number: MYA-3666) was grown in 2X YPAD media with 1.5% agar (w/v) and 1 M *D*-sorbitol. Yeast cultures were incubated at 30°C, with liquid cultures placed on a shaker set to 225 rpm. Agar plates were sealed with parafilm to reduce desiccation. Yeast media were prepared as previously described [24].

*Escherichia coli* (EPI300, Epicenter) was grown at 37°C in Luria Broth (LB) supplemented with antibiotics as needed (50 *µ*g/mL kanamycin). *E. coli* ECGE101 (Δ*dapA*) [25], was grown at 37°C in LB supplemented with 60 *µ*g/mL diaminopimelic acid (DAP), and the appropriate antibiotics (50 *µ*g/mL kanamycin and/or 40 *µ*g/mL gentamycin). Cultures were placed in an incubator set to 37°C, with liquid cultures shaking at 225 rpm.

### Whole-genome sequencing and phylogenetic analysis

Cultures of *C. jejuni* RM1164 and HC1 were grown to saturation, and genomic DNA was extracted using an alkaline lysis protocol as previously described [26]. Whole-genome sequencing was performed by Flow Genomics using their Bacterial Genomes service. Genomic DNA was sequenced on the Oxford Nanopore MinION, with a R10.4.1 flow cell and the SQK-RBK114.96 rapid barcoding kit following manufacturer’s instructions (Oxford Nanopore Technologies). Basecalling was performed on Dorado (v0.9.6), read filtering with Filtlong (v0.3.1), genome assembly with Flye (v2.9.6), polishing with Medaka (v2.2.0), and genome annotation with either Bakta (v1.12) or pLannotate (v1.2.4). Publicly available *C. jejuni* genomes were downloaded from the National Centre for Biotechnology Information (NCBI), and after trimming redundant copies, we were left with 526 unique genomes. In addition, *C. jejuni* genomes of interest (NCTC 11168 and 81-176) were also isolated and included in downstream analyses. Mash (v2.3) was used to generate sketches and distance matrices, and a custom python script converted the matrices into phylogenetic trees. The tree was visualized using the Interactive Tree of Life [27].

### DNA isolation

DNA was isolated from *C. jejuni* and *S. cerevisiae* using a modified alkaline lysis protocol as previously described [26]. We modified step 1 for *C. jejuni* DNA isolation, which is described below. Step 1 was modified for *C. jejuni* to isolate pCJ01 with reduced contamination and at higher concentrations. Step 1: *C. jejuni* transformants that were re-patched twice were grown to saturation on Columbia Blood agar supplemented with sheep blood and the appropriate antibiotics (10 *µ*g/mL vancomycin and 5 *µ*g/mL trimethoprim as background antibiotics, with 30 *µ*g/mL kanamycin for the transformants). Colonies were passaged as large streaks to ensure there were sufficient cells for DNA isolation; typically, a single plate would be used to passage a single colony. Plates were grown over 2-3 days. Cells were then scraped and resuspended in 250 *µ*L resuspension buffer. For *E. coli*, we used the EZ-10 DNA Miniprep Kit to isolate and purify DNA (BioBasic Inc. BS414). DNA was eluted in 30 *µ*L ddH_2_O.

### PCR amplification

All primers are listed in Supplementary S1 Table. Fragments for assembly were amplified using PrimeSTAR GXL DNA Polymerase (Takara, R050A) using the PCR protocol. PCR reactions were run over 30-35 cycles depending on amplification efficiency. A DpnI (New England Biolabs, R0176) was used to removed template DNA from the amplifications, following manufacturer’s protocols. Fragments were run over a 1% agarose gel and purified using the PureLink PCR Purification Kit (Invitrogen, K310001) and eluted using ddH_2_O.

### Screening *C. jejuni* and *E. coli* transformants

Five individual transformant colonies were passaged three times on solid media with the appropriate antibiotic, and DNA was isolated and PCR screened for the inserted kanamycin resistance gene. 1 *µ*L of DNA isolated by alkaline lysis was used as a template in a 20 *µ*L SuperPlex PCR reaction (Takara, 638543). Reactions were performed according to manufacturer’s protocol for 30 cycles.

### Plasmid construction

The pCJ01 plasmid was developed from the pHflu3 plasmid [28] by the addition of the *C. jejuni* origin of replication (*OriC*) and kanamycin resistance gene using yeast assembly. pHflu3 was linearized by XhoI (New England Biolabs, R0146). The *OriC* was amplified from *C. jejuni* RM1164 genomic DNA, and the kanamycin resistance gene was amplified from pMW10 [29] using primers that added 40 bp of overlapping homology to each end (S1 Table). The resulting amplicons and linearized pHflu3 were combined in an equimolar ratio and co-transformed into *S. cerevisiae* spheroplasts, yielding pCJ01. Yeast assembly was conducted as previously described [24] using mixtures of DNA fragments in place of a bacterial donor. Following assembly, DNA was isolated from pooled *S. cerevisiae* transformants and electroporated into *E. coli* EPI300. Individual colonies were screened for correct plasmid assembly by multiplex PCR and diagnostic restriction digestion.

### Electroporation to *C. jejuni*

The preparation of electrocompetent cells was performed as previously described [14]. *C. jejuni* was streaked onto CBA supplemented with 5% sheep blood, 10 *µ*g/mL vancomycin and 5 *µ*g/mL trimethoprim from a frozen glycerol stock and grown for 24 hours at 37°C under microaerophilic conditions. The culture was replated onto a new plate and incubated for an additional 16 hours under the same growth conditions. Following this incubation, the colonies were scraped and resuspended in 2 mL of Brain Heart Infusion (BHI) (Difco^™^211059) liquid broth without antibiotics. Cells were centrifuged at 16,000xg for 5 min at 4°C. The supernatant was discarded, and cell pellets were resuspended in 2 mL of ice-cold wash buffer (63% (v/v) glycerol/37% (w/v) sucrose). The centrifugation and resuspensions were repeated an additional two times. After the final centrifugation, the supernatant was discarded, and the pelleted cells were resuspended in 600 *µ*L of ice-cold wash buffer. The cells were then divided into 50 *µ*L aliquots in sterile microcentrifuge tubes and stored at -80°C.

Electroporation into *C. jejuni* follows as a protocol as previously described [14], with minor modifications. 50 *µ*L of *C. jejuni* cells were combined with 20 *µ*g of pCJ01 in a microcentrifuge tube and incubated on ice for 2-3 minutes. The mix was then transferred to a prechilled 2 mm cuvette (VWR) and pulsed in a GenePulser Xcell (Bio-Rad) at 2.5 kV voltage, 200 Ω resistance, and 25 *µ*F capacitance, with a time constant of ∼5 msec. Cells recovered in 100 *µ*L SOC media (5 mL 2M MgCl_2_, 10 mL 250 mM KCl, 20 mL 1M glucose, 0.5 g/L NaCl, 5 g/L yeast extract, 20 g/L tryptone) on Sheep Blood agar with Columbia Base plate supplemented with 10 *µ*g/mL vancomycin and 5 *µ*g/mL trimethoprim at 37°C under microaerophilic conditions for 5 hours. After recovery, cells were harvested by resuspending in 1 mL BHI broth without antibiotics, followed by centrifugation for 2 min at 14,000xg and discarding the supernatant. Pellets were resuspended in 100 *µ*L of BHI broth and spread across a Sheep Blood agar with Columbia Base plate supplemented with 10 *µ*g/mL vancomycin, 5 *µ*g/mL trimethoprim, and 30 *µ*g/mL kanamycin at 37°C under microaerophilic conditions. Colony forming units could be visualized on plates after 2-3 days.

### Transformation into *E. coli*

Isolated plasmid DNA was added to 50 *µ*L electrocompetent *E. coli* cells in a 1.5 mL tube. The mix was gently pipetted to mix and transferred to a pre-chilled 2 mm electrocuvette. The cuvette was pulsed at 2.5 kV voltage, 200 Ω resistance, and 25 *µ*F capacitance. Cells recovered in 1 mL SOC media at 37°C with shaking at 225 rpm for 1 hour. Cells were collected by centrifugation for 2 min at 10,000xg. The supernatant was discarded, and pellets were resuspended in 100 *µ*L of liquid LB and spread across a 1.5% agar (w/v) LB plate supplemented with or without DAP 60 *µ*g/mL and the appropriate antibiotics (50 *µ*g/mL kanamycin and/or 40 *µ*g/mL gentamycin). Plates were incubated at 37°C, and transformants could be visualized within 24 hours.

### *C. jejuni* plasmid recovery and digests

Following transformation and plating, 10 *C. jejuni* transformants were passaged three times on solid media with the appropriate antibiotics. 5 colonies were harvested from the plates by resuspending in ddH_2_O. Cells were pelleted and DNA was isolated by alkaline lysis. Approximately 1 *µ*L of isolated DNA (∼300 ng) was used to transform into *E. coli* as previously described. For each transformation, plasmid DNA was isolated using the EZ-10 DNA Miniprep Kit (BioBasic Inc. BS414) according to manufacturer’s instructions. Isolated pCJ01 was digested using 5 units of XbaI (New England Biolabs, R0145L) in a 20 *µ*L reaction at 37°C for 1 hour. Complete digestion was confirmed on a 1% agarose gel.

### Conjugation assays from *E. coli* to *C. jejuni*

The *E. coli* ECGE101 Δ*dapA* donor strain [25] harbouring pTA-Mob [30] were grown at 37°C overnight in 5 mL of LB media supplemented with 60 *µ*g/mL DAP, 40 *µ*g/mL gentamycin (donor only), and 50 *µ*g/mL kanamycin. Saturated *E. coli* cultures were diluted 1:10 into 5 mL of fresh LB and grown to an OD_600_ of 0.5. The *C. jejuni* recipient strain was streaked on a Columbia Blood Agar (CBA) plate containing 5% sheep blood and the background antibiotics for 16-20 hours at 37°C under microaerophilic conditions. Cells were then resuspended from the plates in 2 mL of Brain Heart Infusion (BHI) broth without antibiotics and diluted to an OD_600_ of ∼ 1 in 1 mL of BHI media. For the *E. coli* donor or control strain, 1 mL of culture was washed twice by centrifugation at 10,000xg for 5 min at 4°C, discarding the supernatant, and adding 1 mL BHI broth without antibiotics. The *E. coli* donor and control pellets were then resuspended with 1 mL of the *C. jejuni* suspension. The bacterial mix was spun down at 12,000xg for 5 min at 4°C, and resuspended in 100 *µ*L BHI broth without antibiotics. This resuspension was then spread onto CBA plates supplemented with 5% sheep blood and 60 *µ*g/mL DAP. These plates were incubated at 37°C for 5 hours under microaerophilic conditions, and then cells were scraped off with 1 mL of BHI broth, and adjusted final volume of 1 mL in a microfuge tube. A 10-fold serial dilution was performed with BHI broth, and 100 *µ*L of each dilution was plated on selective CBA agar (5% sheep blood, 10 *µ*g/mL vancomycin, 10 *µ*g/mL trimethoprim (TMP), and 30 *µ*g/mL kanamycin). Plates were incubated at 37°C for 2-3 days under microaerophilic conditions.

### Antibiotic susceptibility assays

*C. jejuni* was grown on CBA agar supplemented with 5% sheep blood and the appropriate antibiotic (10 *µ*g/mL vancomycin, 5 *µ*g/mL trimethoprim, and/or 30 *µ*g/mL kanamycin). Cells were scraped and resuspended in 1 mL of BHI broth and diluted to an OD_600_ of 0.5 before being serially diluted in CBA media to either a 10^-4^ or 10^-5^ dilution. 3 *µ*L of each dilution was plated on media containing either 2 *µ*g/mL chloramphenicol, 5 *µ*g/mL gentamycin, 5 *µ*g/mL neomycin, 1 *µ*g/mL tetracycline, 5 *µ*g/mL kanamycin, or on a non-selective control. Plates were incubated for 3 days at 37°C under microaerophilic conditions before images were taken.

## Results

### Whole-genome sequencing of *C. jejuni* strains

Two *C. jejuni* strains were collected from contaminated food and patient-derived samples. Both strains were subject to long-read whole-genome sequencing, revealing genomic characteristics expected from other *C. jejuni* strains [31,32]. Both genomes trend towards the smaller size for *C. jejuni*, with genome sizes of 1.69 Mb and 1.62 Mb for HC1 and RM1164, respectively (Fig 1A/B). Primer pairs designed from *C. jejuni* NCTC 11168 were used to genotype HC1 and RM1164, with expected sizes ranging from 165-650 bp (Fig 1C). Interestingly, sequencing also identified two cryptic plasmids from *C. jejuni* HC1, pTet and pHC1 (Fig 1D/E). pHC1 appears to carry elements of conjugative machinery and virulence factors, although the current annotation seems to suggest that this is an incomplete Type IV Secretion System (T4SS). pTet does not carry complementary conjugation machinery, though it does carry a gene conferring resistance to tetracycline, enabling its continual propagation. No contigs beyond genomic DNA were identified for *C. jejuni* RM1164. A cursory investigation into the evolutionary ancestry of RM1164 and HC1 reveal that HC1 shares a fairly recent common ancestor with the first sequenced *C. jejuni* genome, the flagship strain NCTC 11168 (Fig 1F) [33]. However, RM1164 appears distant, sharing a recent common ancestor with neither HC1, NCTC 11168, nor *C. jejuni* 81-176, another highly virulent isolate of interest [34].

**Fig 1.**
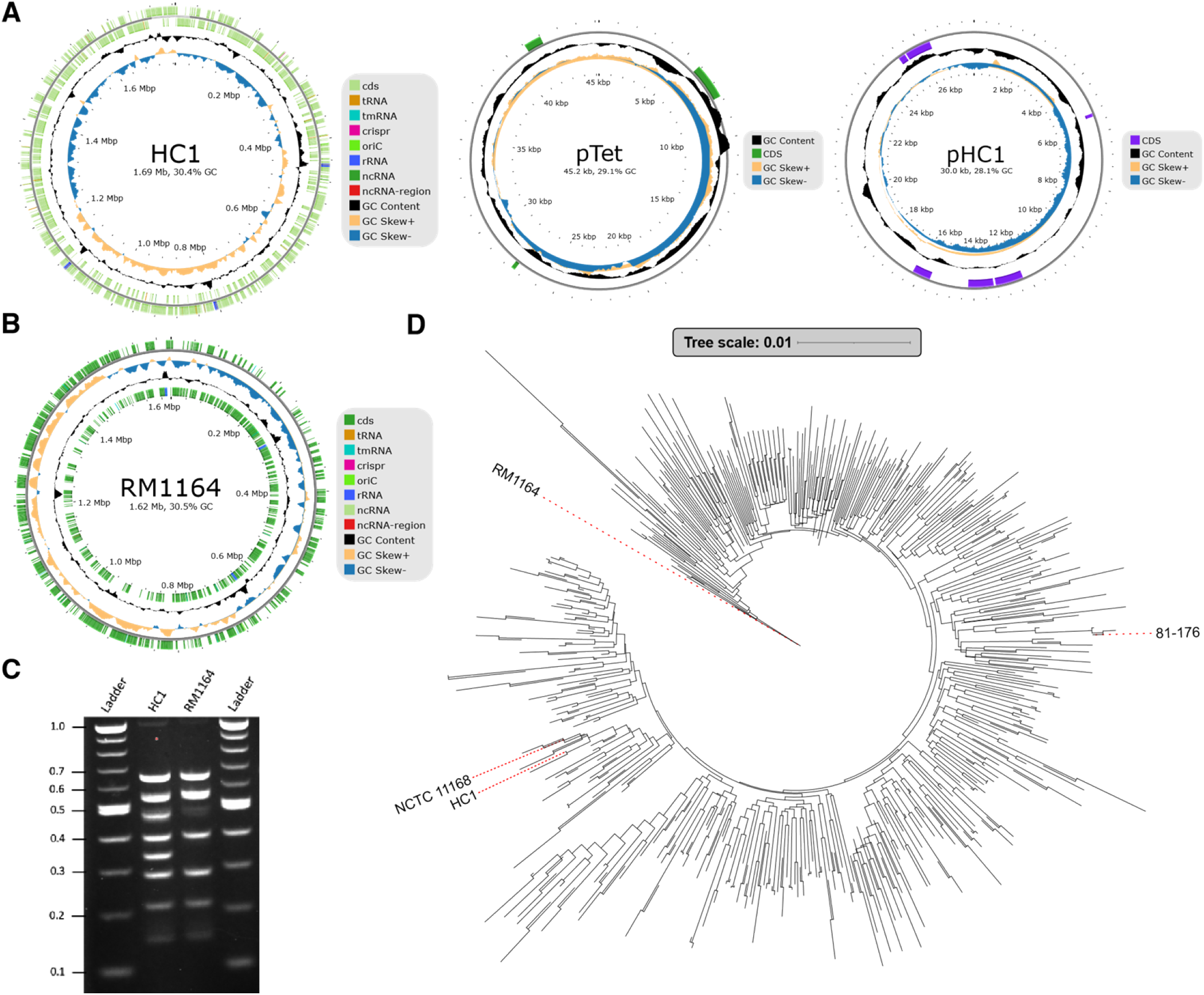
Whole genome sequencing of two patient and sample derived strains of *C. jejuni*. (A) Annotated genome of *C. jejuni* strain HC1. *C. jejuni* strain HC1 also carries two cryptic plasmids, pHC1 which carries conjugative machinery and virulence factors, and pTet which carries a tetracycline resistance gene. (B) Annotated genome of *C. jejuni* strain RM1164. (C) Multiplex PCR genotyping of HC1 and RM1164. Eight primer pairs were designed from *C. jejuni* NCTC 11168 and used against HC1 and RM1164. (D) Phylogenetic relationship of the newly sequenced *C. jejuni* strains HC1 and RM1164 to 526 randomly selected publicly available *C. jejuni* genomes, as well as the reference strains NCTC 11168 and 81-176, based on whole-genome sequence similarity.

### Antibiotic susceptibility assays

To identify appropriate selection conditions for *C. jejuni* RM1164, the response of the strain to a range of common selection markers was assessed via spot plating, comparing growth on selective media to non-selective growth conditions (Fig 2). *C. jejuni* RM1164 demonstrated moderate resistance to lower concentrations of chloramphenicol (2 *µ*g/mL), however higher concentrations (3 *µ*g/mL) were capable of suppressing growth. This strain was highly sensitive to all tested concentrations of gentamycin, neomycin, kanamycin, and tetracycline (Fig 1E). This established at least 3-4 potential selection markers capable of positive selection for each of these strains.

**Fig 2.**
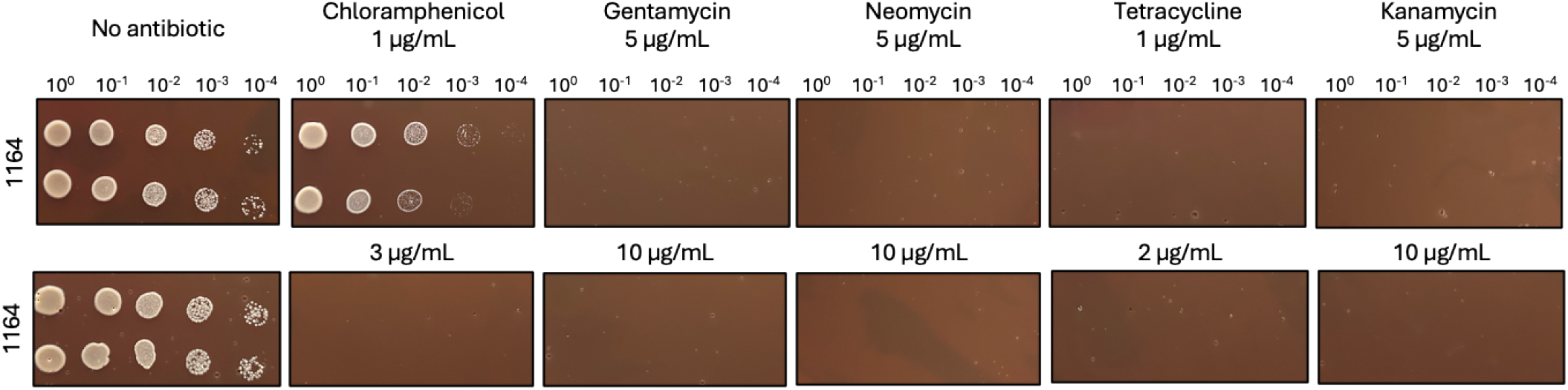
Antibiotic susceptibility assays for wild type *C. jejuni* RM1164. *C. jejuni* RM1164 colonies were passaged, and spot plated onto common antibiotic selection markers. Antibiotic concentrations are measured in *µ*g/mL. A single colony was spotted in duplicate, at dilutions from 10^-1^ to 10^-4^, and photos were obtained after 3 days.

### Developing a shuttle plasmid for *C. jejuni*

We previously designed a series of multi-host plasmids carrying antibiotic selection markers, from which we designed new shuttle plasmids for *C. jejuni* [28]. One such plasmid, pHflu3, carries genes conferring resistance to chloramphenicol, and could be capable of selection in alternative hosts by cloning in *C. jejuni* codon-optimized kanamycin resistance genes. To ensure replication in *C. jejuni*, the putative *OriC* was identified and cloned onto the plasmid, resulting in pCJ01 (Fig 3A).

**Fig 3.**
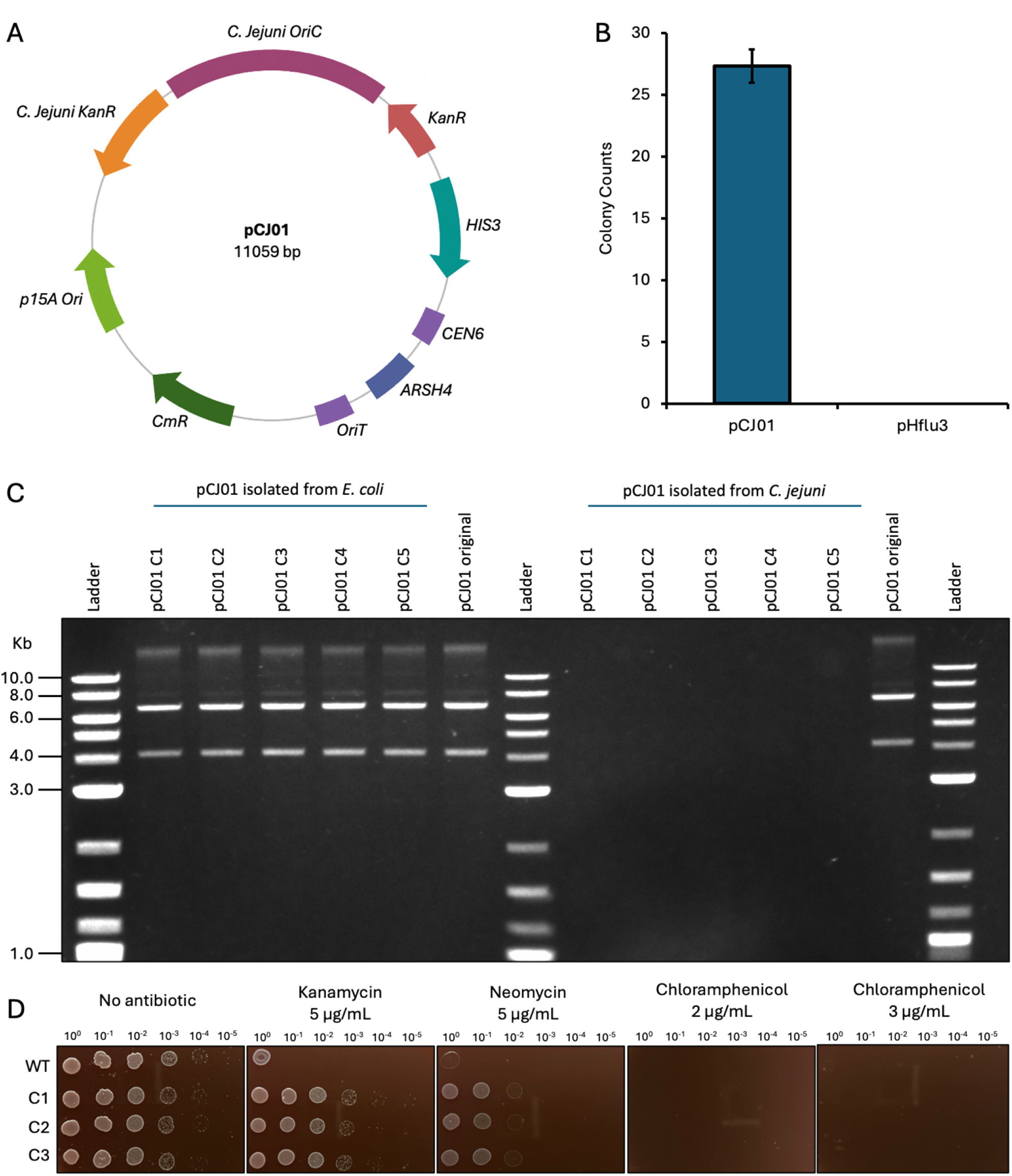
Developing the pCJ01 shuttle plasmid for *C. jejuni*. (A) Plasmid map for pCJ01. Kan^R^, kanamycin resistance gene; p15a Ori, plasmid origin of replication; Cm^R^, chloramphenicol resistance gene; OriT, plasmid origin of transfer; ARSH4, yeast autonomous replicating sequence; CEN6, yeast centromere DNA element; HIS3, yeast selection marker; OriC, *C. jejuni* origin of replication. Maps were created with BioRender.com. (B) Colony counts of *C. jejuni* following transformation of pCJ01 and pHflu3 (negative control). Bars represent an average colony count from 3 independent experiments. Error bars represent the standard error from the mean. (C) Restriction digest with XbaI of pCJ01 recovered from transformants of *E. coli* and *C. jejuni*. pCJ01 original were the plasmids used for the *C. jejuni* transformation. (D) Antibiotic susceptibility assays for *C. jejuni* transformants harbouring pCJ01 and no plasmid *C. jejuni* (WT). Cultures were spotted at dilutions from 10^-1^ to 10^-5^, and images were taken after 3 days.

With the prior antibiotic susceptibility assays demonstrating sensitivity to kanamycin for RM1164, successful plasmid replication of pCJ01 in *C. jejuni* was observed via growth on selective media with kanamycin and neomycin (Fig 3B/D). To verify that resistance was conferred by transformation of pCJ01, *C. jejuni* and *E. coli* transformants were screened by restriction digest, with clear and expected band sizes appearing from the *E. coli* prep (Fig 3C). No bands could be observed from *C. jejuni* transformants, consistent with both a lower copy number of the plasmid in *C. jejuni* and poor recovery owing to insufficient domestication.

### Strain-specific conjugation into *C. jejuni*

Following the successful demonstration of pCJ01 transformation into *C. jejuni* and *E. coli*, we sought to determine if pCJ01 could be transferred from a non-pathogenic *E. coli* strain into *C. jejuni*. Establishing a functional bacterial conjugation system into *C. jejuni* is expected to be a tremendous step forward for *Campylobacter* genetic engineering, enabling the transfer of high molecular weight payloads beyond what is capable by standard transformation [35]. Similar advancements have been transformative for other relatively undomesticated microorganisms [26]. As pCJ01 carries a plasmid origin of transfer (*OriT*), it can be mobilized by a helper plasmid through a process known as conjugation in *trans* [30,36]. Conjugation from an auxotrophic *E. coli* donor strain carrying the helper plasmid and either pCJ01 or pHflu3 demonstrated functional *trans*-conjugation of pCJ01 into *C. jejuni* (Fig 4A). pHflu3 can be considered a negative control due to lacking the *C. jejuni* putative *OriC* and the *C. jejuni* codon-optimized kanamycin resistance gene. To confirm resistance was conferred in *C. jejuni* by pCJ01 conjugation, seven transconjugants were isolated and PCR screened to detect for pCJ01, with all five colonies validated as successful transconjugants (Fig 4B).

**Fig 4.**
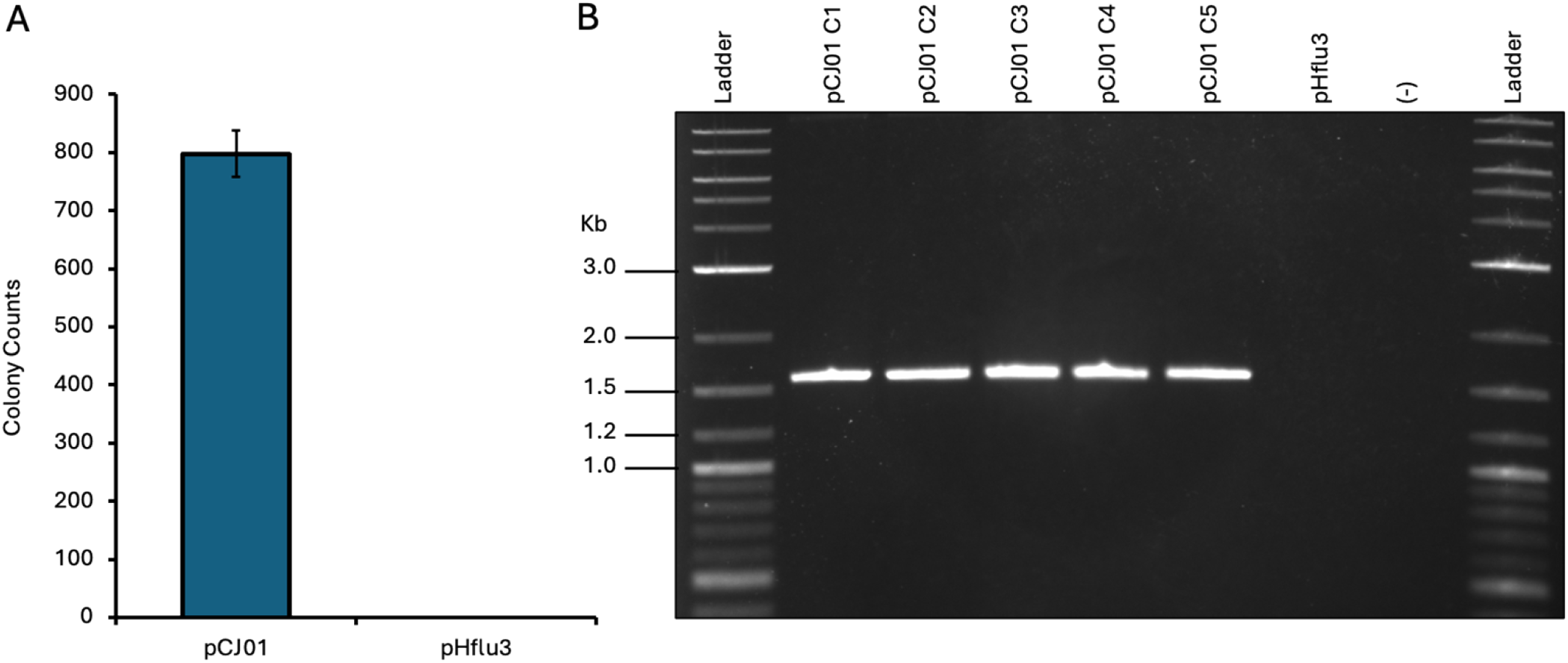
Demonstrating strain-specific plasmid transfer from *E. coli* to *C. jejuni* using conjugation. (A) Colony counts of *C. jejuni* following conjugation with pCJ01 or pHflu3. Bars represent an average colony count from three independent experiments. Error bars represent the standard error from the mean. (B) PCR screen of *C. jejuni* transconjugants of pCJ01. Primers specific to the *C. jejuni* Kan^R^ cassette produced a band of the expected size in pCJ01, but not in pHflu3. (-) represents the no-template control for the PCR reaction. The expected amplicon size is 1,678 bp.

## Discussion

This work expands the genetic toolkit for *C. jejuni* by characterizing two clinically relevant, evolutionarily distinct isolates. Whole-genome sequencing revealed strain-specific differences between *C. jejuni* HC1 and RM1164, including two cryptic plasmids in *C. jejuni* HC1. However, no plasmids were detected in *C. jejuni* strain RM1164, consistent with reports that plasmid content varies widely among *C. jejuni* isolates [37–39]. In addition, including evolutionarily diverse *C. jejuni* strains is important because genetic tools often show strain-dependent performance and can be difficult to apply across different isolates [40,41]. We also tested antibiotic susceptibility in *C. jejuni* RM1164 and identified several selection markers that were useful for subsequent work in this strain.

An important result of this study is the development of a mobilizable shuttle plasmid and a conjugation mediated delivery system for *C. jejuni* RM1164. Previous genetic work in *C. jejuni* has relied heavily on natural transformation, which is often strain dependent [14,40,42]. In this study, we demonstrate conjugation-based plasmid delivery from *E. coli* donors into *C. jejuni* RM1164, providing an alternative route for DNA transfer. Unlike natural transformation, conjugation does not rely on the natural competence of recipient cells [43].

Beyond technical development, the strains and plasmids described here have practical implications for *C. jejuni* detection. *C. jejuni* and other clinically relevant strains can be difficult to culture and manipulate genetically, which has limited the design and testing of engineered biosensors for contaminated food products. A system that supports plasmid transfer and maintenance in *C. jejuni* could be useful for future development of improved detection tools for this pathogen. Current detection methods are often culture-based and can take time, which limits how quickly contamination can be identified. Making clinically relevant *C. jejuni* strains more accessible to genetic manipulation may help support the design and testing of new assay strategies. However, further work is needed to quantify conjugation efficiencies across a broader range of clinical isolates and to determine how reliably this system performs under conditions relevant to food processing. In the longer term, tools like this could help support earlier monitoring in food production settings and improve food safety.

## Author contributions

V.N.: conceptualization, formal analysis, investigation, methodology, writing - original draft, writing - review and editing; G.W.F.: formal analysis, investigation, methodology, writing - review and editing; T.J.: investigation, writing - review and editing; A.E.: conceptualization, investigation; M.J.R.: conceptualization, methodology, writing - review and editing; B.J.K. conceptualization, formal analysis, funding acquisition, methodology, resources, supervision, writing - review and editing.

## Competing interests statement

The authors declare there are no competing interests.

## Data availability

BAM files for whole-genome sequencing of *C. jejuni* RM1164 and HC1 were deposited in the Sequence Read Archive under accession number: PRJNA1438426.

### Acknowledgment

We thank Dr. Carole Creuzenet for providing pMW10 plasmid, which was used as the source of the kanamycin resistance gene in this study.

## Funding statement

This research was funded by an NSERC Alliance Grant 548857-2019 awarded to M.J.R. In addition, this research was supported by the Natural Sciences and Engineering Research Council of Canada (NSERC), grant number: RGPIN-2025-05428 awarded to B.J.K.

## Notes

### Competing Interest Statement

The authors have declared no competing interest.

## References

1. Delahoy MJ, Shah HJ, Weller DL, Ray LC, Smith K, McGuire S, et al. Preliminary Incidence and Trends of Infections Caused by Pathogens Transmitted Commonly Through Food — Foodborne Diseases Active Surveillance Network, 10 U.S. Sites, 2022. MMWR Morb Mortal Wkly Rep. 2023;72: 701–706. doi:10.15585/mmwr.mm7226a1

2. Tack DM, Ray L, Griffin PM, Cieslak PR, Dunn J, Rissman T, et al. Preliminary Incidence and Trends of Infections with Pathogens Transmitted Commonly Through Food — Foodborne Diseases Active Surveillance Network, 10 U.S. Sites, 2016–2019. MMWR Morb Mortal Wkly Rep. 2020;69: 509–514. doi:10.15585/mmwr.mm6917a1

3. Drudge C, Greco S, Kim J, Copes R. Estimated Annual Deaths, Hospitalizations, and Emergency Department and Physician Office Visits from Foodborne Illness in Ontario. Foodborne Pathog Dis. 2019;16: 173–179. doi:10.1089/fpd.2018.2545

4. Kaakoush NO, Castaño-Rodríguez N, Mitchell HM, Man SM. Global Epidemiology of Campylobacter Infection. Clin Microbiol Rev. 2015;28: 687–720. doi:10.1128/CMR.00006-15

5. Igwaran A, Okoh AI. Human campylobacteriosis: A public health concern of global importance. Heliyon. 2019;5: e02814. doi:10.1016/j.heliyon.2019.e02814

6. Facciolà A, Riso R, Avventuroso E, Visalli G, Delia SA, Laganà P. Campylobacter: from microbiology to prevention. J Prev Med Hyg. 2017;58: E79–E92. doi:10.15167/2421-4248/jpmh2017.58.2.672

7. Man SM. The clinical importance of emerging Campylobacter species. Nat Rev Gastroenterol Hepatol. 2011;8: 669–685. doi:10.1038/nrgastro.2011.191

8. Dietrich J, Hammerl J-A, Johne A, Kappenstein O, Loeffler C, Nöckler K, et al. Impact of climate change on foodborne infections and intoxications. J Health Monit. 2023;8: 78–92. doi:10.25646/11403

9. Takamiya M, Ozen A, Rasmussen M, Alter T, Gilbert T, Ussery DW, et al. Genome Sequence of Campylobacter jejuni strain 327, a strain isolated from a turkey slaughterhouse. Stand Genomic Sci. 2011;4: 113–122. doi:10.4056/sigs.1313504

10. Ocejo M, Oporto B, Lavín JL, Hurtado A. Whole genome-based characterisation of antimicrobial resistance and genetic diversity in Campylobacter jejuni and Campylobacter coli from ruminants. Sci Rep. 2021;11: 8998. doi:10.1038/s41598-021-88318-0

11. Djeghout B, Bloomfield SJ, Rudder S, Elumogo N, Mather AE, Wain J, et al. Comparative genomics of Campylobacter jejuni from clinical campylobacteriosis stool specimens. Gut Pathog. 2022;14: 45. doi:10.1186/s13099-022-00520-1

12. Garcia-Fernandez A, Janowicz A, Marotta F, Napoleoni M, Arena S, Primavilla S, et al. Antibiotic resistance, plasmids, and virulence-associated markers in human strains of Campylobacter jejuni and Campylobacter coli isolated in Italy. Front Microbiol. 2024;14: 1293666. doi:10.3389/fmicb.2023.1293666

13. Tikhomirova A, McNabb ER, Petterlin L, Bellamy GL, Lin KH, Santoso CA, et al. Campylobacter jejuni virulence factors: update on emerging issues and trends. J Biomed Sci. 2024;31: 45. doi:10.1186/s12929-024-01033-6

14. Davis L, Young K, DiRita V. Genetic Manipulation of Campylobacter jejuni: Epsilon Proteobacteria. Curr Protoc Microbiol. 2008;10: 8A.2.1-8A.2.17. doi:10.1002/9780471729259.mc08a02s10

15. Golz JC, Stingl K. Natural Competence and Horizontal Gene Transfer in Campylobacter. In: Backert S, editor. Fighting Campylobacter Infections. Cham: Springer International Publishing; 2021. pp. 265–292. doi:10.1007/978-3-030-65481-8_10

16. Korkus J, Sałata P, Thompson SA, Paluch E, Bania J, Wałecka-Zacharska E. The role of cydB gene in the biofilm formation by Campylobacter jejuni. Sci Rep. 2024;14: 26574. doi:10.1038/s41598-024-77556-7

17. Yao R, Alm RA, Trust TJ, Guerry P. Construction of new Campylobacter cloning vectors and a new mutational cat cassette. Gene. 1993;130: 127–130. doi:10.1016/0378-1119(93)90355-7

18. Golden NJ, Camilli A, Acheson DWK. Random Transposon Mutagenesis of Campylobacter jejuni. O’Brien AD, editor. Infect Immun. 2000;68: 5450–5453. doi:10.1128/IAI.68.9.5450-5453.2000

19. Mandal RK, Jiang T, Kwon YM. Essential genome of Campylobacter jejuni. BMC Genomics. 2017;18: 616. doi:10.1186/s12864-017-4032-8

20. Teh AHT, Lee SM, Dykes GA. Identification of potential Campylobacter jejuni genes involved in biofilm formation by EZ-Tn5 Transposome mutagenesis. BMC Res Notes. 2017;10: 182. doi:10.1186/s13104-017-2504-1

21. Wilson DL, Bell JA, Young VB, Wilder SR, Mansfield LS, Linz JE. Variation of the natural transformation frequency of Campylobacter jejuni in liquid shake culture. Microbiology. 2003;149: 3603–3615. doi:10.1099/mic.0.26531-0

22. Holt JP, Grant AJ, Coward C, Maskell DJ, Quinlan JJ. Identification of Cj1051c as a Major Determinant for the Restriction Barrier of Campylobacter jejuni Strain NCTC11168. Appl Environ Microbiol. 2012;78: 7841–7848. doi:10.1128/AEM.01799-12

23. Johnson JG, Carpentier S, Spurbeck RR, Sandhu SK, DiRita VJ. Genome Sequences of Campylobacter jejuni 81–176 Variants with Enhanced Fitness Relative to the Parental Strain in the Chicken Gastrointestinal Tract. Genome Announc. 2014;2: e00006–14. doi:10.1128/genomeA.00006-14

24. Karas BJ, Jablanovic J, Irvine E, Sun L, Ma L, Weyman PD, et al. Transferring whole genomes from bacteria to yeast spheroplasts using entire bacterial cells to reduce DNA shearing. Nat Protoc. 2014;9: 743–750. doi:10.1038/nprot.2014.045

25. Brumwell SL, MacLeod MR, Huang T, Cochrane RR, Meaney RS, Zamani M, et al. Designer Sinorhizobium meliloti strains and multi-functional vectors enable direct inter-kingdom DNA transfer. Martinez-Abarca F, editor. PLOS ONE. 2019;14: e0206781. doi:10.1371/journal.pone.0206781

26. Karas BJ, Diner RE, Lefebvre SC, McQuaid J, Phillips APR, Noddings CM, et al. Designer diatom episomes delivered by bacterial conjugation. Nat Commun. 2015;6: 6925. doi:10.1038/ncomms7925

27. Letunic I, Bork P. Interactive Tree of Life (iTOL) v6: recent updates to the phylogenetic tree display and annotation tool. Nucleic Acids Res. 2024;52: W78–W82. doi:10.1093/nar/gkae268

28. Hamadache S, Huang YK, Shedeed A, Syed A, Karas BJ. Deletion of HindIIR and HindIIIR improves DNA transfer via electroporation to Haemophilus influenzae Rd. The Microbe. 2024;4: 100125. doi:10.1016/j.microb.2024.100125

29. Wösten MMSM, Boeve M, Koot MGA, Van Nuenen AC, Van Der Zeijst BAM. Identification of Campylobacter jejuni Promoter Sequences. J Bacteriol. 1998;180: 594–599. doi:10.1128/JB.180.3.594-599.1998

30. Cochrane RR, Shrestha A, Severo De Almeida MM, Agyare-Tabbi M, Brumwell SL, Hamadache S, et al. Superior Conjugative Plasmids Delivered by Bacteria to Diverse Fungi. BioDesign Res. 2022;2022: 9802168. doi:10.34133/2022/9802168

31. Duong T, Konkel ME. Comparative studies of Campylobacter jejuni genomic diversity reveal the importance of core and dispensable genes in the biology of this enigmatic food-borne pathogen. Curr Opin Biotechnol. 2009;20: 158–165. doi:10.1016/j.copbio.2009.03.004

32. Shyaka A, Kusumoto A, Asakura H, Kawamoto K. Whole-Genome Sequences of Eight Campylobacter jejuni Isolates from Wild Birds. Genome Announc. 2015;3: e00315–15. doi:10.1128/genomeA.00315-15

33. Parkhill J, Wren BW, Mungall K, Ketley JM, Churcher C, Basham D, et al. The genome sequence of the food-borne pathogen Campylobacter jejuni reveals hypervariable sequences. Nature. 2000;403: 665–668. doi:10.1038/35001088

34. Poly F, Threadgill D, Stintzi A. Genomic Diversity in Campylobacter jejuni : Identification of C. jejuni 81-176-Specific Genes. J Clin Microbiol. 2005;43: 2330–2338. doi:10.1128/JCM.43.5.2330-2338.2005

35. Smillie C, Garcillán-Barcia MP, Francia MV, Rocha EPC, De La Cruz F. Mobility of Plasmids. Microbiol Mol Biol Rev. 2010;74: 434–452. doi:10.1128/MMBR.00020-10

36. Brumwell SL, Van Belois KD, Giguere DJ, Edgell DR, Karas BJ. Conjugation-Based Genome Engineering in Deinococcus radiodurans. ACS Synth Biol. 2022;11: 1068–1076. doi:10.1021/acssynbio.1c00524

37. Marasini D, Fakhr M. Exploring PFGE for Detecting Large Plasmids in Campylobacter jejuni and Campylobacter coli Isolated from Various Retail Meats. Pathogens. 2014;3: 833–844. doi:10.3390/pathogens3040833

38. Morita D, Arai H, Isobe J, Maenishi E, Kumagai T, Maruyama F, et al. Whole-Genome and Plasmid Comparative Analysis of Campylobacter jejuni from Human Patients in Toyama, Japan, from 2015 to 2019. Gao B, editor. Microbiol Spectr. 2023;11: e02659–22. doi:10.1128/spectrum.02659-22

39. Bacon DJ, Alm RA, Burr DH, Hu L, Kopecko DJ, Ewing CP, et al. Involvement of a Plasmid in Virulence of Campylobacter jejuni 81-176. Barbieri JT, editor. Infect Immun. 2000;68: 4384–4390. doi:10.1128/IAI.68.8.4384-4390.2000

40. Beauchamp JM, Leveque RM, Dawid S, DiRita VJ. Methylation-dependent DNA discrimination in natural transformation of Campylobacter jejuni. Proc Natl Acad Sci. 2017;114. doi:10.1073/pnas.1703331114

41. Yamamoto S, Iyoda S, Ohnishi M. Stabilizing Genetically Unstable Simple Sequence Repeats in the Campylobacter jejuni Genome by Multiplex Genome Editing: a Reliable Approach for Delineating Multiple Phase-Variable Genes. Parkhill J, editor. mBio. 2021;12: e01401–21. doi:10.1128/mBio.01401-21

42. Wiesner RS, Hendrixson DR, DiRita VJ. Natural Transformation of Campylobacter jejuni Requires Components of a Type II Secretion System. J Bacteriol. 2003;185: 5408–5418. doi:10.1128/JB.185.18.5408-5418.2003

43. Zeng X, Ardeshna D, Lin J. Heat Shock-Enhanced Conjugation Efficiency in Standard Campylobacter jejuni Strains. Elkins CA, editor. Appl Environ Microbiol. 2015;81: 4546–4552. doi:10.1128/AEM.00346-15

